# WeSeqMiner: A Weka package for building machine-learning models for sequence data

**DOI:** 10.1101/217802

**Authors:** Daniel Hogan, Bharathikumar V. Maruthachalam, C. Ronald Geyer, Anthony Kusalik

## Abstract

In most cases, the application of machine learning techniques to biological sequence data requires a vector representation of the sequences. Extracting the numerical features from sequence data can be time consuming, especially if the user lacks programming skills. To this end, we propose a Weka package called WeSeqMiner, which provides several useful filters for extracting numerical features from sequence data for use in the Weka machine learning workbench. Motivated with an example, we show that the WeSeqMiner package integrates well with the Weka API, allowing transformations to be incorporated into Weka workflows for predictive model generation. WeSeqMiner can be installed by pointing the Weka package manager to the URL github.com/djhogan/WeSeqMiner/raw/master/WeSeqMiner.zip. The Javadoc for WeSeqMiner classes can be accessed at djhogan.github.io/seqminer.

## 1 Introduction

A number of high-throughput technologies generate large datasets relating peptide sequences to a biological property of interest. For example, in antibody phage display these peptides are variants of a complementarity determining region (CDR), often CDRH3, which is the primary determinant of antibody specificity [2]. The property of interest is the target-specificity of the antibodies incorporating these CDRs. Another technology is peptide microarrays, which contain several thousand spots of unique peptides in a grid-like arrangement on a glass surface. These peptides may be potential epitopes as in the case of epitope discovery, or predicted kinase targets as in the case of kinome analysis.

The large volume of data produced by high-throughput technologies like antibody phage display and peptide microarrays makes machine learning a viable option for discovering functions relating sequence to biological state, function, or condition. Bioinformatics is replete with such examples [4]. Unfortunately, the majority of machine learning techniques cannot be directly applied to sequence data due to its string representation, and the way to vectorize these strings is not obvious. Researchers must resort to extracting features from the string data that retain the salient information or signal for their prediction task. One feature extraction strategy is to represent each amino acid in a sequence as a vector of numerical properties. Two such strategies are to use columns of a substitution matrix (e.g. BLOSUM62) or the Kidera Factors. The Kidera Factors [3] are the 10 principal components uncovered from a principal component analysis of 188 physico-chemical properties of amino acids; therefore, each of the 20 amino acids would be represented by a unique vector of length 10 in this principal component space.

To simplify sequence feature extraction for users of the Weka machine learning workbench [8] we propose a Weka package called WeSeqMiner, which conforms to the Weka API, and thus benefits from the Weka ecosystem, providing a convenient interface for feature extraction and integration with upstream tasks, like reading data, and downstream tasks like feature selection or classifier/regressor training.

## 2 Software Features

WeSeqMiner provides Weka with 6 new filters that extract numerical attributes (features) from a basic string attribute representing a protein sequence. Three filters (*AminoAcidCounts*, *DipeptideCounts*, and *KideraFactors*) summarize the content of a sequence but retain no positional information. Each of these three filters accepts an option flag (-R) that, when turned on, normalizes the extracted features by the length of the sequence, *n* (or *n* − 1 in the case of *DipeptideCounts*). *OneHotEncoding* and *BlosumEncoding* encode each amino acid (position) in the sequence as a vector. *OneHotEncoding* achieves this by setting one of twenty elements in each vector to unity to denote the amino acid in the sequence. *BlosumEncoding* encodes the amino acid by its row in the BLOSUM62 matrix. Because the dimensionality of the output for these two filters is a function of the sequence length, the input sequences must be of equal length. Variable length sequences can be made equal in length either by trimming or by applying the *SlidingWindow* filter, the sixth filter provided by WeSeqMiner. It creates overlapping sequences of equal length from a larger sequence. Because WeSeqMiner conforms to the Weka API, it benefits from many of the features offered by the Weka ecosystem, including knowledgeflow pipelines.

## 3 Case Study

Phage display selections were carried out using a well-characterized single-framework synthetic antibody library named Library-F [1, 5]. Five rounds of solid-phase panning were conducted against seven recombinant protein targets (full-length extracellular domains of Notch1, Notch2, Notch3, Jagged1, Jagged2, Axl and Mer) as described previously [6]. After each selection round, the CDRH3 region was amplified from the phage selection output and subjected to Ion Torrent next-generation sequencing [7] on an Ion Personal Genome Machine according to manufacturer's instructions. Following Ion Torrent sequencing, base-called reads were parsed by barcodes, quality trimmed, and filtered. Filtered sequences were translated, identified, and counted. A Weka knowledgeflow was constructed to assess the relative effectiveness of features extracted by WeSeqMiner classes for predicting the enrichment of CDRH3 sequences during the antibody selection process [2] against any of the seven targets.

The knowledgeflow incorporated five analyses. Analyses 1-3 used each of the feature sets produced by the filters *AminoAcidCounts*, *DipeptideCounts*, and *KideraFactors* with a random forest (RF) classifier. A RF classifier was selected because it (1) is capable of discovering complex relationships between predictor and outcome variables, (2) naturally resists overfitting, and (3) requires minimal configuration. The 4th analysis tested the union of features produced by the filters with an RF classifier. The 5th analysis required no feature extraction, accepting the raw sequence as input and using a string kernel SVM. The dataset used in the analysis was a balanced collection of both sequences that were not observed after 5 rounds of panning (”negatives”), and sequences that were still present after 5 rounds of panning (”positives”). Ten-fold cross-validation was used to evaluate the accuracy of each classifier resulting from the training.

The accuracies of the five trained classifiers were 84.4% (all features + RF), 82.0% (amino acid counts + RF), 83.9% (dipeptide counts + RF), 77.8% (Kidera factors + RF), and 82.0% (raw sequence + SVC). For the four RF classifiers, the threshold of votes was varied to determine the receiver operating characteristics (ROCs). The ROC curves are shown in Figure 1. With the exception of performance visualizations, which were generated using Python, the pipeline was built and executed entirely within the Weka workbench, obviating any need to write program code.

**Figure 1:**
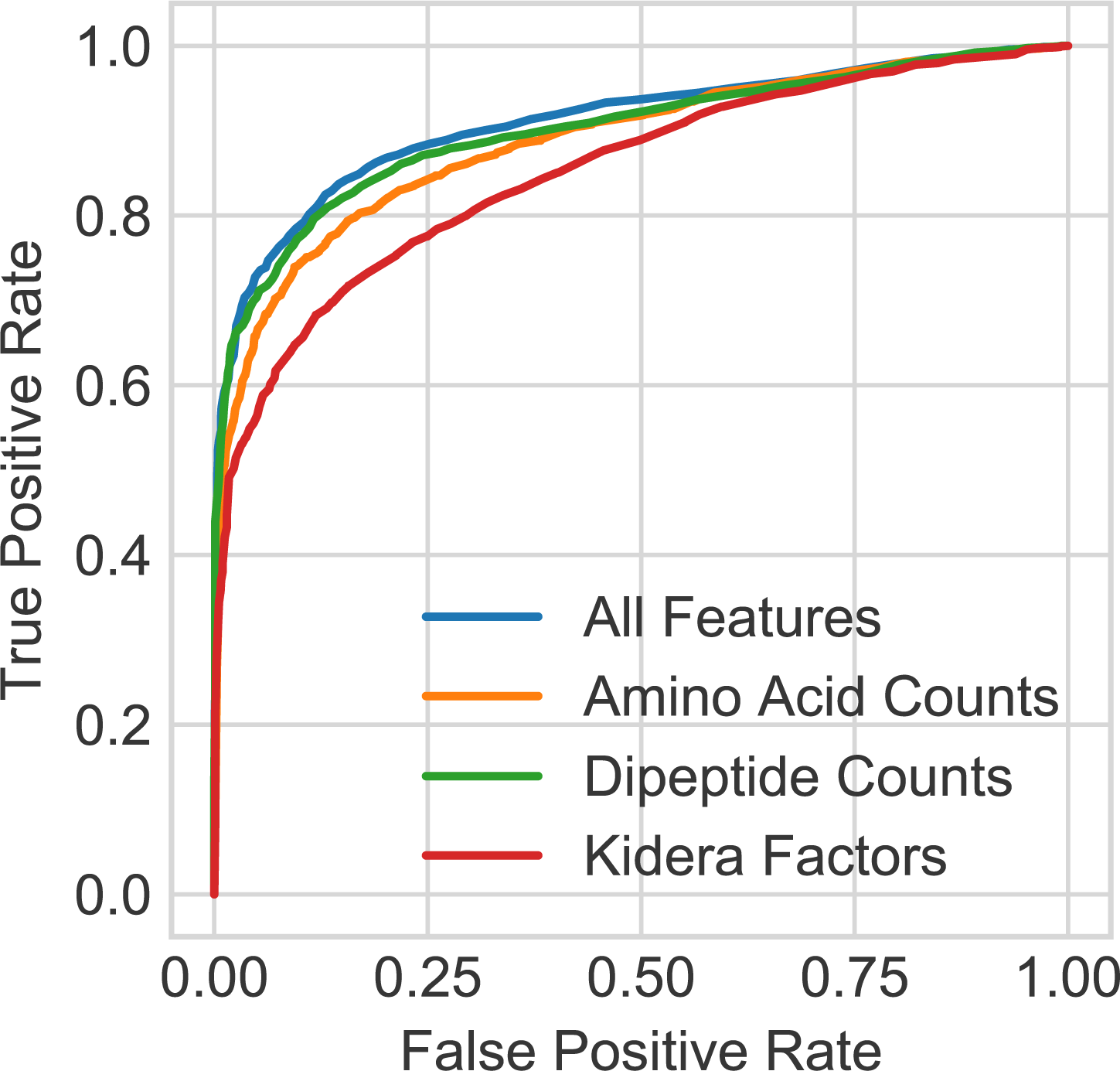
ROC for RF classifiers trained on different WeSeqMiner feature sets.

## 4 Conclusion

Our case study showed that WeSeqMiner adds to the Weka ecosystem a convenient method for extracting features from sequence data, such that models can be built, trained, and tested on this feature data using Weka. With the help of WeSeqMiner it was shown that four feature sets are comparable in their ability to predict clone enrichment, with the combination of the three feature sets performing the best. It was also shown that there are patterns in Library-F CDRH3 sequences that can predict the enrichment against any of the seven different targets. WeSeqMiner could be used to carry out similar analyses in other settings. For example, WeSeqMiner could be used with data from epitope arrays, where one is interested in relating amino acid sequence properties to the flourescence of the peptides on the array. Such a relationship would be of interest to researchers because this flourescence indicates the presence of bound serum antibodies; therefore, the relationship could be used to screen peptides during array design to maximize the number of epitopes placed on the array.

